# LOBSTERS: a modular vector series with a diverse set of Cas nucleases and plasmid selection markers for multiplex genome editing in *Saccharomyces cerevisiae*

**DOI:** 10.1101/2025.10.05.680590

**Authors:** Satoshi Okada, Emiko Kusumoto, Goro Doi, Shitomi Nakagawa, Takashi Ito

## Abstract

The budding yeast *Saccharomyces cerevisiae* is a central model organism in genetics and synthetic biology, yet efficient multiplex genome editing remains difficult because many toolkits are restricted by limited plasmid selection markers and reduced efficiency when targeting multiple loci. In our previous work, we introduced a CRISPR-based system incorporating three nucleases with distinct PAM specificities but it was available only with the *URA3* marker. Here we present LOBSTERS, an expanded modular vector series that retains the PAM-diverse nucleases (SpCas9, SaCas9, and enAsCas12a) while extending marker options to seven. This design enables the simultaneous use of multiple plasmids in one transformation, supporting scalable and flexible genome editing. Proof-of-concept experiments demonstrated efficient *ADE2* and *ADE3* deletions with colorimetric readouts, coordinated tagging of essential proteins (Cdc3 and Cse4) without compromising function, and recapitulation of three quantitative trait variants (*RME1, TAO3*, and *MKT1*) underlying sporulation efficiency. Together these results establish LOBSTERS as a robust and versatile platform for multiplex genome editing in *S. cerevisiae*. By enabling coordinated modification of essential proteins, genetic interactions, and quantitative trait variants, LOBSTERS provides a broadly applicable resource for functional cell biology and synthetic biology in yeast.

**Significance Statement:**

- LOBSTERS integrates three PAM-diverse nucleases with seven plasmid selection markers, overcoming the single-marker limitation of previous yeast genome editing toolkits.
- The system enables efficient simultaneous editing at multiple loci, demonstrated by functional tagging of essential proteins and recapitulation of quantitative trait variants.
- By broadening the editable space and lowering barriers to complex genotype construction, LOBSTERS provides a widely applicable resource for yeast cell biology and synthetic biology.

## INTRODUCTION

The budding yeast *Saccharomyces cerevisiae* has long been an important model organism in genetics and synthetic biology, owing to its tractable genetics and efficient homologous recombination. The emergence of CRISPR-Cas technologies has further accelerated yeast genome engineering, enabling precise and programmable manipulations. Numerous CRISPR-based toolkits have since been developed, aiming to streamline workflows and expand the scale of achievable modifications (DiCarlo *et al*., 2013; Laughery *et al*., 2015; Generoso *et al*., 2016; Jessop-Fabre *et al*., 2016; Świat *et al*., 2017; Degreif *et al*., 2018; Verwaal *et al*., 2018).

One major class of systems is designed for integration at predefined “safe harbor” loci, which provide stable and predictable expression environments. EasyClone-MarkerFree exemplifies this approach, enabling markerless integration of expression cassettes with high efficiency and reproducibility but limiting edits to specific genomic locations (Jessop-Fabre *et al*., 2016). Similarly, the Multiplex Yeast Toolkit (MYT) combines Golden Gate cloning with CRISPR-Cas9 to support the simultaneous integration of multiple expression cassettes at validated sites (Shaw *et al*., 2023). These systems are powerful for pathway engineering when stable expression at safe loci is desired, but their applicability is constrained to predetermined genomic regions.

A complementary class of toolkits supports editing at arbitrary genomic loci, broadening flexibility but often at the cost of efficiency or marker versatility. The pCAS system and related vectors allow single-vector expression of Cas9 and single guide RNA (sgRNA) for general gene disruption or tagging, though with limited selection markers (Ryan *et al*., 2014; Laughery *et al*., 2015). More recent systems, including CMI (Meng *et al*., 2023), MULTI-SCULPT (Gong *et al*., 2022), and CREEPY (Zhao *et al*., 2023), enhance efficiency for multiplex editing or episomal manipulation, yet challenges remain when scaling to multiple loci simultaneously. Thus, while arbitrary locus–targeting approaches expand the editable space, they are still constrained by the availability of selection markers and by declining efficiency as the number of edits increases.

In our previous work, we developed a CRISPR-based toolkit that incorporated three Cas nucleases with distinct protospacer adjacent motif (PAM) specificities, thereby expanding the proportion of the genome that could be targeted, but relied on a single selection marker, *URA3* (Okada *et al*., 2021). Building on this foundation, we now present LOBSTERS (League Of Backbone plasmid vector Series To Expand the Range of Selection markers), a modular vector series that retains the PAM-diverse nucleases (SpCas9, SaCas9, and enAsCas12a) (Jinek *et al*., 2012; Ran *et al*., 2015; Kleinstiver *et al*., 2019) while expanding the selection markers to seven. This design both maintains broad genomic editability and enables the simultaneous use of multiple plasmids in a single transformation, thereby providing a flexible and scalable platform for multiplex genome engineering in yeast.

Importantly, by facilitating multi-locus editing in a single transformation, LOBSTERS provides an experimental framework well suited for modern cell biology. It supports simultaneous interrogation of essential proteins, pathway components, and regulatory variants, thereby offering opportunities to probe genetic networks and emergent phenotypes that cannot be addressed by sequential editing approaches.

## RESULTS

### Expansion of the backbone vector series for yeast genome editing

In a previous report, we constructed a series of backbone vector plasmids for genome editing in the budding yeast *S. cerevisiae* (Okada *et al*., 2021). The vector series was designed to meet three essential requirements: (1) both Cas protein and sgRNA/crRNA are encoded on a single plasmid, (2) expression of Cas protein and/or sgRNA/crRNA can be artificially induced via the well-characterized *GAL1* promoter, and (3) target sequences of sgRNA/crRNA can be easily incorporated using the Golden Gate Assembly method (Engler *et al*., 2008). These vectors were initially constructed using three CRISPR-Cas systems: SpCas9 from *Streptococcus pyogenes* (PAM = NGG), SaCas9 from *Staphylococcus aureus* (PAM = NNGRRT), and enAsCas12a from *Acidaminococcus* sp. (PAM = TTYN, VTTV, and TRTV). By incorporating nucleases with distinct PAM specificities, this toolkit expanded the editable space of the yeast genome, allowing access to a broader range of target sites compared to single-nuclease systems. This PAM diversity remains a central strength of the platform, as it minimizes the risk of “uneditable” loci and provides users with greater flexibility in experimental design (Okada *et al*., 2021).

Although functional, the vectors in this series had a major limitation: they were available only with the *URA3* marker. This restricted their use to *ura3* mutant strains, even though researchers often need to modify strains with diverse genotypes, including those constructed using the *URA3* marker. Moreover, targeting multiple loci under this design required either laborious sequential editing with different nucleases, which could exploit the broadened target space, or single-step editing with one nuclease and multiple sgRNAs/crRNAs, which was restricted to a limited target space. To overcome these limitations, we expanded the range of selection markers for the backbone vector series (Figure 1). The vector architecture retains the RNA processing elements from our previous system (Okada et al., 2021). For SpCas9 and SaCas9, sgRNAs are flanked by hammerhead and hepatitis delta virus (HDV) ribozymes at the 5′ and 3′ ends, respectively; self-cleavage at both termini generates mature sgRNAs with precisely defined ends. For the enAsCas12a system, a tRNA(Gly) sequence positioned upstream of the crRNA exploits endogenous tRNA processing (Zhang *et al*., 2019): RNase P cleaves at the tRNA 3′ end to release the crRNA 5′ terminus, while HDV ribozyme processes the 3′ end, together producing functional crRNAs with correct terminal structures. For each of the three CRISPR-Cas systems (SpCas9, SaCas9, and enAsCas12a), seven selection markers (*HIS3, TRP1, LEU2, URA3, KanMX, HphMX*, and *NatMX*) are now available. These markers include both auxotrophic and drug-resistance options, thereby making this series applicable to a broad spectrum of commonly used yeast strains. Moreover, by retaining the nuclease diversity of the original system while greatly expanding marker availability, this series enables full exploitation of the broadened editable space, even in multiplex editing with a single transformation. The expanded series was named LOBSTERS (<u>L</u>eague <u>O</u>f <u>B</u>ackbone plasmid vector <u>S</u>eries <u>T</u>o <u>E</u>xpand the <u>R</u>ange of <u>S</u>election markers for genome editing in budding yeast). This design provides greater flexibility for users working with diverse strain backgrounds and paves the way for complex editing strategies, including multiplex gene disruptions, tagging, and reconstruction of polygenic traits.

**Figure 1.**
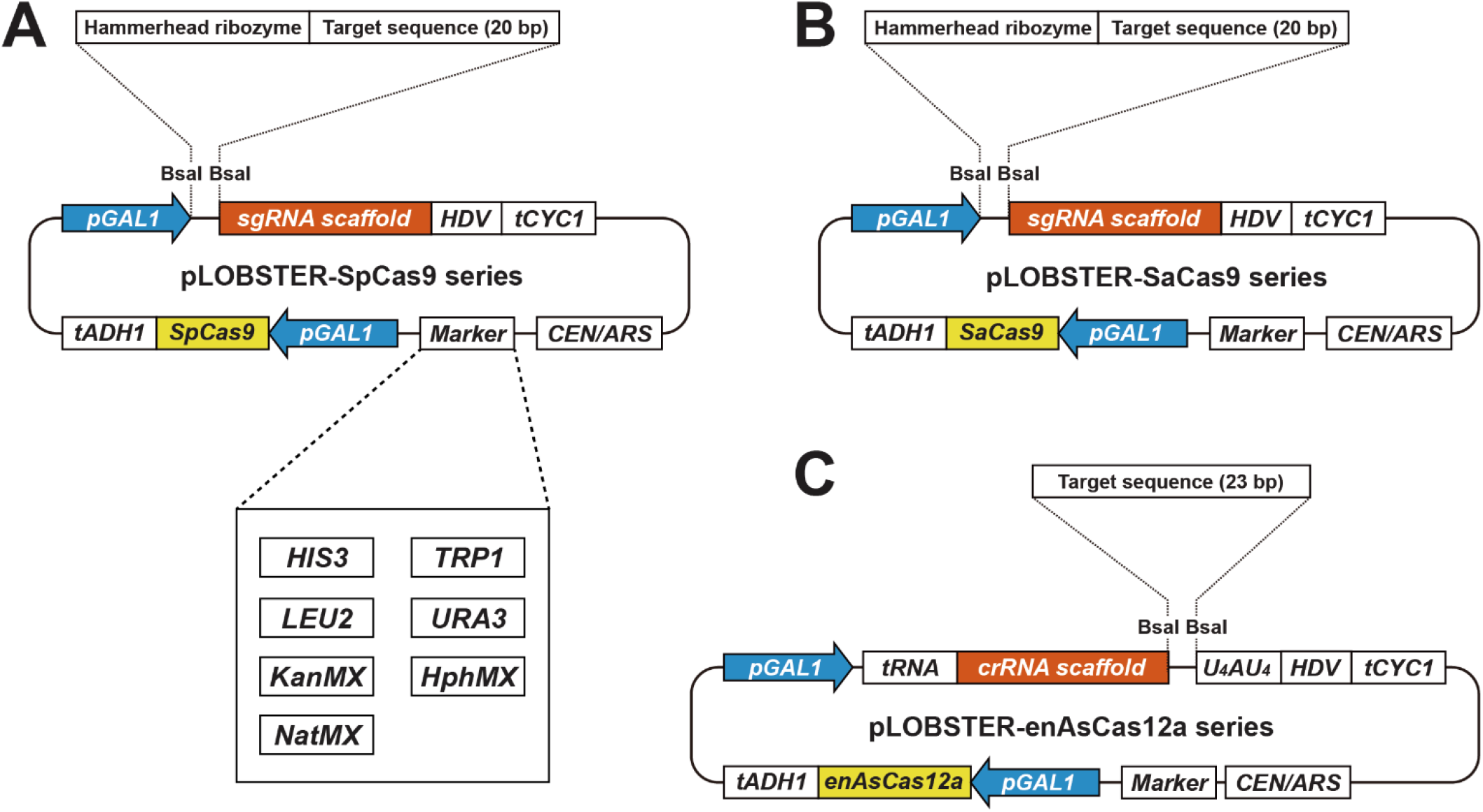
Schematic representation of LOBSTERS. (A) pLOBSTER-SpCas9 series. (B) pLOBSTER-SaCas9 series. (C) pLOBSTER-enAsCas12a series. *pGAL1, GAL1* promoter; *tADH1, ADH1* terminator; *tCYC1, CYC1* terminator; *HDV*, HDV ribozyme; *tRNA*, glycine tRNA; *U*_*4*_*AU*_*4*_, 9-mer encoding “UUUUAUUUU” for improvement of genome editing efficiency; *CEN/ARS*, centromere and autonomously replicating sequence; *HIS3, HIS3* marker cassette; *TRP1, TRP1* marker cassette; *LEU2, LEU2* marker cassette; *URA3, URA3* marker cassette; *KanMX*, G418 resistance marker cassette; *HphMX*, hygromycin B resistance marker cassette; *NatMX*, nourseothricin (clonNAT) resistance marker cassette.

### Validation across multiple markers and Cas systems

To rigorously evaluate the versatility of LOBSTERS across distinct Cas nucleases and a broadened set of selection markers, we carried out complete open reading frame (ORF) deletions at well-characterized loci. As test cases, we chose *ADE2* and *ADE3*, whose loss-of-function phenotypes yield unambiguous colony color changes, making them particularly useful for rapid and reliable phenotypic scoring.

*ADE2* encodes phosphoribosylaminoimidazole carboxylase, an enzyme in the adenine biosynthetic pathway (Hieter *et al*., 1985). Its deletion leads to the accumulation of a red-colored intermediate, producing colonies that appear dark red on adenine-limited medium. *ADE3* encodes C1-tetrahydrofolate synthase, another enzyme required for adenine biosynthesis (McKenzie and Jones, 1977). In an *ade2*Δ background, additional deletion of *ADE3* prevents the accumulation of the red pigment, causing colonies to shift from red to white (Koshland *et al*., 1985). This hierarchical phenotype provides a convenient assay for sequential gene disruption and has long served as a benchmark in yeast genetics.

For each of the three CRISPR-Cas systems in LOBSTERS—SpCas9, SaCas9, and enAsCas12a—we designed target sites within the *ADE2* and *ADE3* ORFs (Figure S1A–F) and constructed corresponding editing plasmids with different selection markers. Donor fragments were generated by PCR, each carrying 100 bp of flanking homology (50 bp upstream and 50 bp downstream of the ORF). Transformants were selected on media corresponding to the relevant markers, enabling us to test the full range of vector backbones.

Editing efficiency was first assessed for *ADE2* by quantifying the proportion of red colonies (Figure 2, A and B). Across diverse marker–Cas combinations, efficiency ranged from 51.7% to 85.3%, indicating that all backbones are competent for efficient editing. For *ADE3*, deletions were performed in an *ade2*Δ background, and efficiency was measured as the percentage of white colonies (Figure 2, C and D). These values ranged from 51.3% to 86.1%, consistent across nucleases and markers. By contrast, transformation with control plasmids lacking Cas and gRNA expression produced no change in colony color, confirming that the observed phenotypes were Cas dependent. Colony PCR of a subset of clones verified complete deletion of the targeted ORFs (data not shown).

**Figure 2.**
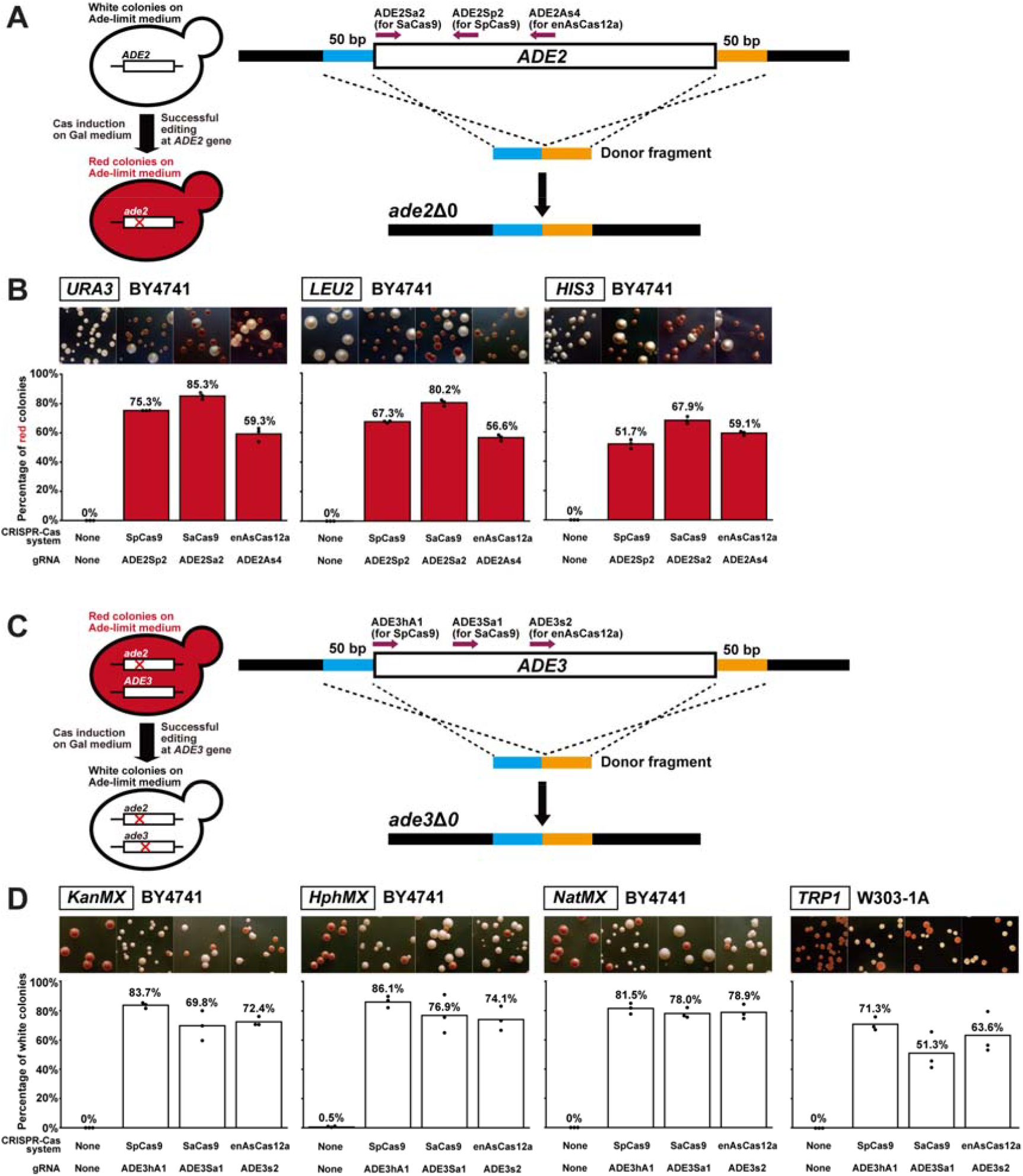
Phenotypic validation of ORF disruption using LOBSTERS. (A) Schematic representation of deletion of the entire *ADE2* ORF and conversion of colony color. The dark magenta arrows represent the positions of the target sequences for each of the three CRISPR-Cas systems. The 50-bp regions used as the 5′- and 3′-homology arms of the 100-bp donor PCR fragment are colored in blue and orange, respectively (schematic not proportional to actual size). The *ade2*Δ cells accumulate red pigments on an adenine-limited medium. (B) Representative colony images after transformation of the genome editing plasmids with the donor PCR products on adenine-limited galactose-containing medium (top) and the efficiency of colony color conversion (bottom). Marker genes for the genome editing plasmids are shown on top of the plate images. Each red bar indicates the percentage of red colonies in each experimental condition shown at the bottom. The value is the average of 3 biological replicates indicated by black dots. (C) Schematic representation of deletion of the entire *ADE3* ORF and conversion of colony color. The dark magenta arrows represent the positions of the target sequences for each of the three CRISPR-Cas systems. The 50-bp regions used as the 5′- and 3′-homology arms of the 100-bp donor PCR fragment are colored in blue and orange, respectively. While *ade2*Δ cells accumulate red pigments on an adenine-limited medium, *ade2*Δ *ade3*Δ cells fail to do so and form white colonies. (D) Representative colony images after transformation of the genome editing plasmids with the donor PCR products on adenine-limited galactose-containing medium (top) and the efficiency of colony color conversion (bottom). Marker genes for the genome editing plasmids are shown on top of the plate images. Each white bar indicates the percentage of white colonies in each experimental condition shown at the bottom. The value is the average of 3 biological replicates indicated by black dots. Note that *TRP1* marker was tested using W303-1A (*trp1-1*) because BY4741 strains carry the wild-type *TRP1* allele. For (B) and (D), complete ORF deletion was confirmed by colony PCR in a subset of clones (data not shown).

Together, these results provide strong validation that every backbone in the LOBSTERS series can support genome editing with robust efficiency, regardless of the Cas nuclease or marker used. The use of visually scorable colorimetric phenotypes not only enabled straightforward quantification of editing success but also highlighted the practical advantages of simple genetic readouts for benchmarking multiplex genome editing systems in yeast. This demonstration establishes a firm foundation for applying LOBSTERS to more complex editing scenarios, where multiple markers and nucleases must be coordinated in parallel.

### Simultaneous double knock-in enabled by marker diversity

A defining advantage of the LOBSTERS vector series is its inclusion of multiple selection markers, which permits concurrent use of distinct genome editing plasmids in a single transformation. Earlier single-marker systems prevented co-introduction of multiple constructs; LOBSTERS overcomes this limitation and enables coordinated multi-locus editing. To demonstrate the potential, we performed a proof-of-concept experiment inserting fluorescent proteins into two essential genes with well-characterized localization properties, *CDC3* and *CSE4*.

*CDC3* encodes a septin protein that assembles into a ring structure at the bud neck, forming a scaffold for cytokinesis (Caviston *et al*., 2003). C-terminal tagging disrupts its localization and causes morphological defects (Huh *et al*., 2003; Dubreuil *et al*., 2019), whereas insertion into the N-terminal region predicted to be devoid of secondary structure preserves normal function. We therefore used SpCas9 to insert an infrared fluorescent protein miRFP682 (Matlashov *et al*., 2020) into the N-terminal loop, enabling visualization of the septin ring without compromising growth (Figure 3, A and B and Figure S1G).

**Figure 3.**
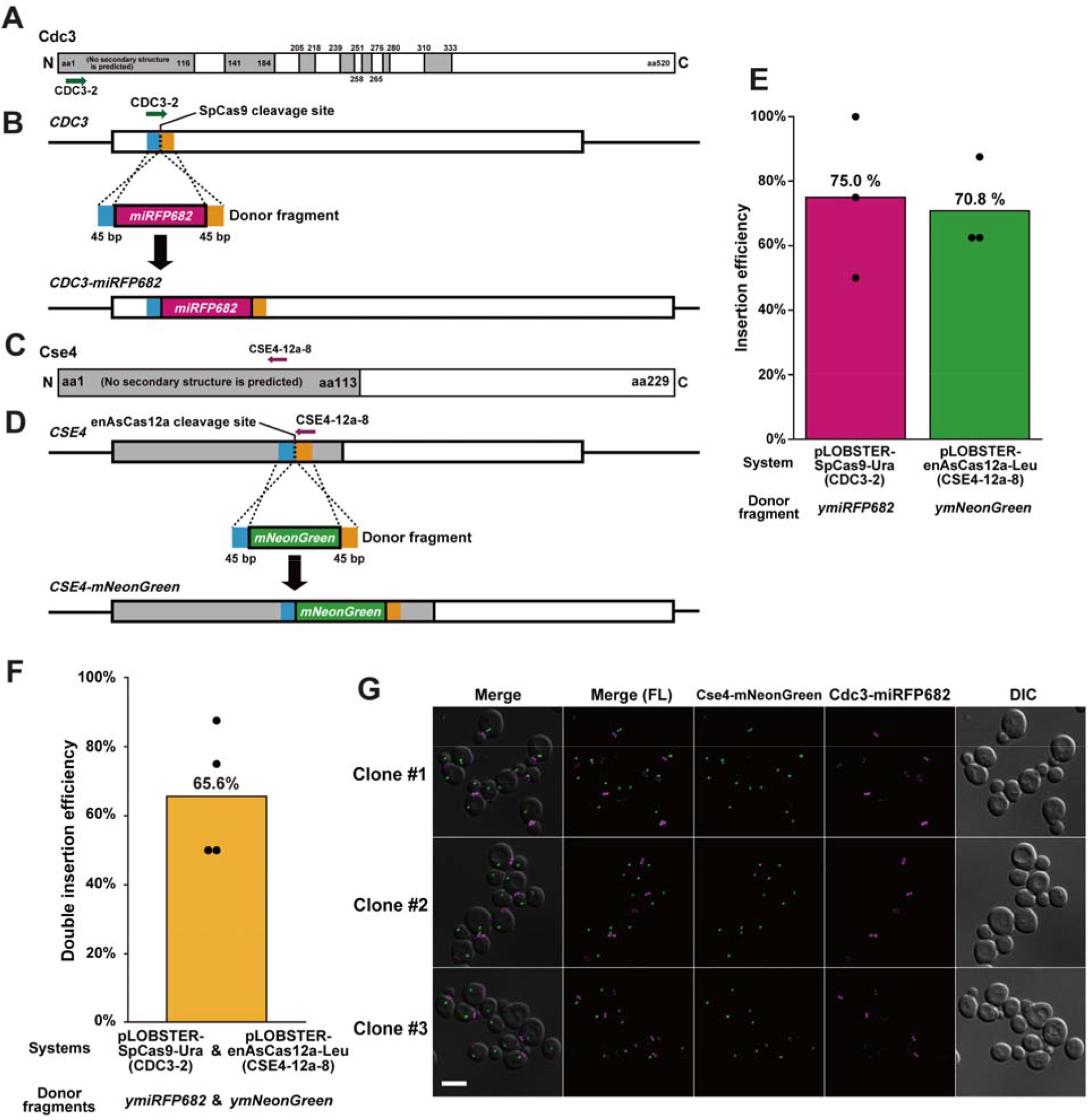
Simultaneous double gene fragment insertion into essential genes using a combination of two pLOBSTER plasmids. (A) Domain structure of the Cdc3 protein. Gray rectangles indicate regions in which no secondary structure is predicted. (B) The *miRFP682* gene fragment insertion process by genome editing using the SpCas9 system at a target sequence indicated by an arrow colored green. The 45-bp regions used as the 5′- and 3′-homology arms are colored blue and orange, respectively. (C) Domain structure of the Cse4 protein. A gray rectangle indicates the region in which no secondary structure is predicted. (D) The *mNeonGreen* gene fragment insertion process by genome editing using the enAsCas12a system at a target sequence indicated by an arrow colored dark magenta. The 45-bp regions used as the 5′- and 3′-homology arms are colored blue and orange, respectively. (E) The efficiency of the single genome editing using a CRISPR-Cas system and a donor fragment. The combinations of a CRISPR-Cas system and a donor fragment are shown below the graph. Bars indicate the average insertion efficiency over three experiments (n = 24 in total). Black dots show the insertion efficiency of each experiment (n = 8 for each). Successful genome editing was confirmed by colony PCR, which showed an increase in PCR product size corresponding to the inserted fluorescent protein gene. (F) The efficiency of the double genome editing using two CRISPR-Cas systems and two donor fragments simultaneously. The combinations of CRISPR-Cas systems and donor fragments are shown below the graph. The bar indicates the average insertion efficiency over four experiments (n = 32 in total). Black dots show the insertion efficiency of each experiment (n = 8 for each). Successful genome editing was confirmed by colony PCR. (G) Representative images of the double-genome-edited *CDC3-miRFP682 CSE4-mNeonGreen* cells. Each row shows an image series of an independent clone. Images are composed of the superimposition of mNeonGreen fluorescent images (green), miRFP682 fluorescent images (magenta), and DIC images (grayscale). Scale bar, 5 μm.

*CSE4* encodes the centromere-specific histone H3 variant essential for kinetochore assembly. C-terminal fusions impair activity and cause temperature sensitivity (Wisniewski *et al*., 2014), but N-terminal loop insertions are tolerated (Zhou *et al*., 2011; Yan *et al*., 2019). Guided by this, we used enAsCas12a to insert a green-yellow fluorescent protein mNeonGreen (Shaner *et al*., 2013) into the N-terminal region, producing a functional fluorescent fusion (Figure 3, C and D and Figure S1H). Both single knock-ins were efficient (70.8–75.0%, Figure 3E). When both plasmids and donor fragments were co-transformed, double knock-ins of *CDC3*-miRFP682 and *CSE4*-mNeonGreen were obtained at 65.6% efficiency (Figure 3F), only modestly lower than single edits. Fluorescence microscopy confirmed correct localization of both proteins, and cells harboring both fusions retained normal morphology (Figure 3G). These outcomes illustrate that simultaneous tagging of two essential proteins is feasible with high precision, without compromising their native functions.

Together, these results demonstrate that LOBSTERS supports simultaneous editing of essential loci at high efficiency and accuracy. By combining marker diversity with nuclease versatility, the system provides a practical and scalable strategy for parallel modifications. This capability not only simplifies experimental workflows but also opens opportunities for interrogating cellular processes that depend on multiple loci, expanding the experimental possibilities of yeast genome engineering. Such marker-assisted multi-locus strategies may also be applied to diverse experimental contexts in yeast genetics and synthetic biology.

### Simultaneous triple-locus editing for quantitative trait reconstruction

The laboratory strain S288c and its derivatives are well known for their extremely low sporulation efficiency, in sharp contrast to the robust sporulation phenotype of the SK1 strain. This difference has long been used as a model for dissecting the genetic basis of quantitative traits in yeast. Pioneering work by Deutschbauer and Davis (2005) identified three quantitative trait variants (QTVs) in SK1—RME1(ins-308A), TAO3(E1493Q), and MKT1(D30G)—as major determinants of this phenotypic divergence. When these three SK1 alleles are introduced into the S288c background, sporulation efficiency is substantially restored, demonstrating that the combined effect of only a few nucleotides can drastically alter a complex biological process. These findings established a classic example of how naturally occurring polymorphisms contribute to phenotypic variation and provided a benchmark for evaluating new genome editing technologies.

To assess the capacity of LOBSTERS for simultaneous editing of multiple loci, we attempted to reconstruct this well-characterized quantitative trait in the S288c background. Using BY4741 and BY4742 as host strains, we designed donor fragments for each QTV (Figure S1I-K) and employed the corresponding Cas systems to introduce the three nucleotide substitutions. A schematic overview of this strategy is presented in Figure 4A. Consistent with previous editing efforts, the efficiency of single knock-ins was high, ranging from 79.2% to 91.7% at individual loci (Figure 4B). When all three edits were attempted in the same transformation, triple knock-in efficiencies of 37.5% in BY4741 and 29.2% in BY4742 were obtained (Figure 4C), confirming that multiplex editing across several loci is feasible, albeit with lower efficiency than single-locus editing.

**Figure 4.**
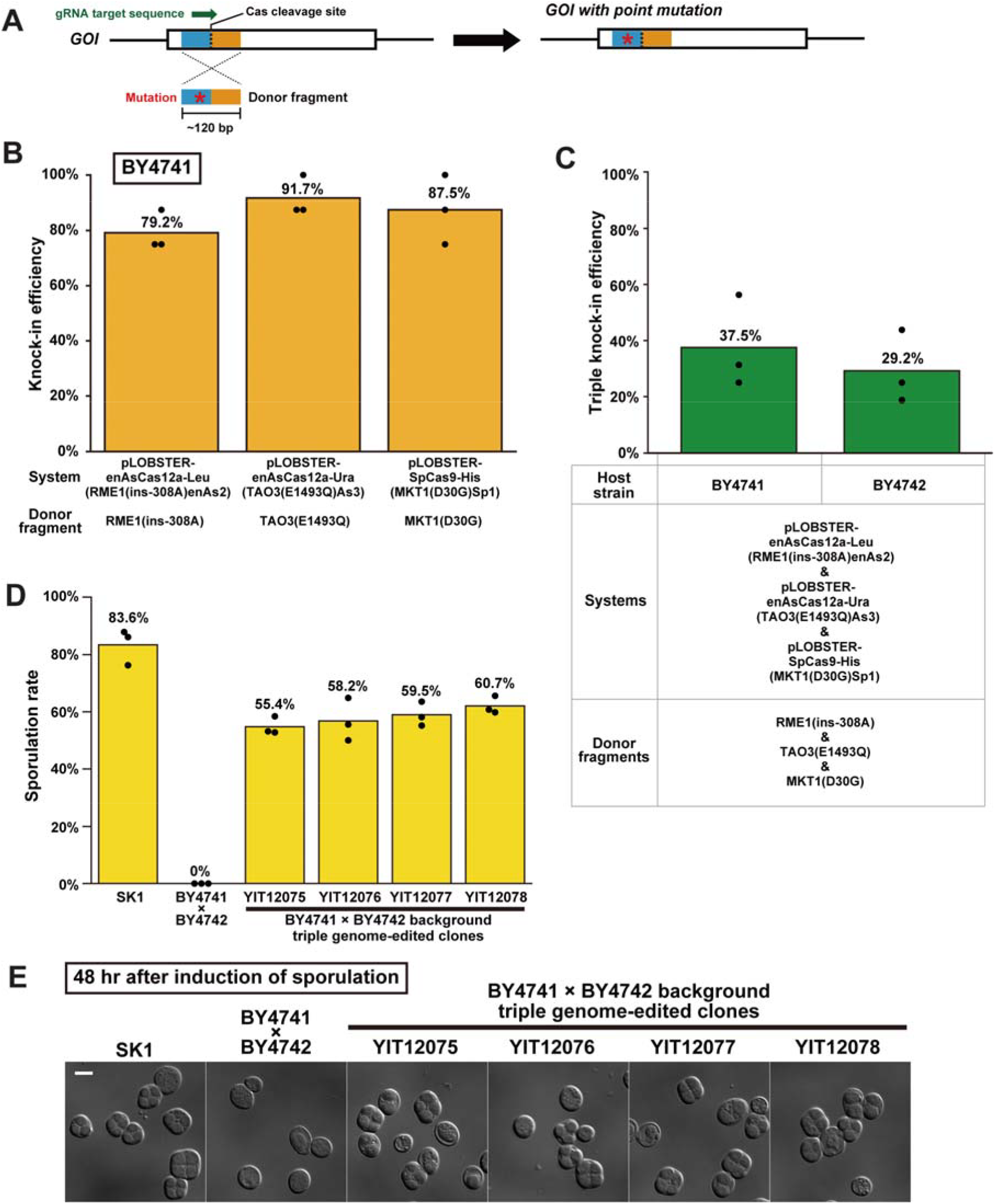
Simultaneous triple genome editing using a combination of three pLOBSTER plasmids. (A) Schematic representation of introducing a single nucleotide/amino acid substitution in the product of a gene of interest (GOI) using CRISPR-Cas systems. An arrow colored green indicates the position of the target sequence. The 5’- and 3’-homology arms are colored blue and orange, respectively. A red asterisk indicates the point mutation in the donor fragment. (B) The knock-in efficiency of the single substitutions into each of the three genes using a CRISPR-Cas system in BY4741 strain. Orange bars indicate the average knock-in efficiency over three experiments (n = 24 in total). Black dots show the knock-in efficiency of each experiment (n = 8 for each). Successful genome editing was confirmed by colony PCR followed by Sanger sequencing of the PCR products. (C) The triple knock-in efficiency of substitutions into the three genes using CRISPR-Cas systems. Green bars indicate the average knock-in efficiency over three experiments (n = 96 in total). Black dots show the knock-in efficiency of each experiment (n = 32 for each). (D) Sporulation efficiency of the SK1 strain, the parental strain in this study (BY4741 x BY4742), and the genome-edited strains (YIT12075, YIT12076, YIT12077, and YIT12078). Each strain was assayed for sporulation efficiency 48 hours after induction of sporulation. (E) Representative DIC images of the strains used in (D). The cells were fixed using 70% ethanol. Scale bar, 5 μm.

Phenotypic assays validated the functional consequences of these edits. As expected, SK1 displayed high sporulation efficiency (83.6%), whereas the BY4741 × BY4742 diploid control strain failed to sporulate within 48 h (0%). In contrast, strains carrying all three SK1-derived alleles exhibited markedly improved sporulation efficiencies, reaching 55.4–60.7% after 48 h (Figure 4, D and E). These values closely mirror those reported by Deutschbauer and Davis (2005), who demonstrated that the introduction of these three QTVs into S288c reconstituted approximately 60% of the sporulation efficiency phenotype observed in SK1.

Taken together, these results illustrate that LOBSTERS not only supports efficient multi-locus genome editing but also enables the faithful reconstruction of naturally occurring quantitative trait variants. By recapitulating a classic genotype–phenotype relationship in yeast, this work highlights the utility of LOBSTERS for dissecting the genetic architecture of complex traits and for validating causal polymorphisms identified through quantitative genetics.

## DISCUSSION

The LOBSTERS vector series addresses key limitations of existing yeast genome editing toolkits by combining nuclease diversity with an expanded repertoire of selection markers. Toolkits designed for safe harbor integration provide stable and predictable expression but are limited to predefined loci, while those enabling arbitrary locus targeting broaden flexibility yet are often constrained by marker availability and reduced efficiency in multiplex contexts. LOBSTERS bridges this gap by retaining three PAM-diverse nucleases and extending the choice of selection markers to seven, thereby supporting both a wide editable space and combinatorial editing strategies within a single transformation.

Our proof-of-principle experiments underscore the versatility of the system. Robust *ADE2* and *ADE3* deletions validated the efficiency of individual vectors across marker–Cas combinations (Figure 2), while double knock-ins into essential genes demonstrated that precise insertions can be achieved without loss of native protein function (Figure 3). Moreover, triple-locus editing recapitulated quantitative trait variants underlying sporulation efficiency, highlighting the utility of LOBSTERS in dissecting complex genotype–phenotype relationships (Figure 4). Together, these results show that combining PAM diversity with marker flexibility enables systematic exploration of genetic architecture.

Despite these strengths, reduced efficiency in multi-locus editing compared with single events remains a challenge. Factors such as competition among plasmids, cellular stress from concurrent DNA breaks, and donor fragment uptake may contribute. Approaches including optimization of plasmid copy number, refinement of donor design, or controlled timing of nuclease activity are likely to improve outcomes. Future developments may also incorporate marker recycling strategies to further expand editing capacity.

Beyond its immediate applications in budding yeast, LOBSTERS establishes a design principle that balances nuclease diversity and marker flexibility. In basic genetics, it enables systematic interrogation of gene networks and quantitative trait loci, while in synthetic biology, it supports combinatorial pathway engineering and strain optimization. For cell biologists, the demonstrated capacity to tag essential proteins and reconstruct quantitative traits illustrates how multiplex genome editing can accelerate mechanistic studies. By overcoming the limitations of earlier single-marker or locus-restricted systems, LOBSTERS provides a robust and scalable framework for multiplex genome engineering in *S. cerevisiae*.

## CONCLUSION

The LOBSTERS vector series integrates three PAM-diverse nucleases with seven selection markers to broaden the editable space of the yeast genome. Marker diversity further allows the simultaneous use of multiple plasmids in a single transformation, enabling efficient multi-locus genome editing. This design overcomes the limitations of earlier systems restricted by single markers or predefined loci. Demonstrated by gene deletions, protein tagging, and trait reconstruction, LOBSTERS offers a flexible and scalable framework for multiplex genome engineering in *S. cerevisiae*.

## MATERIALS AND METHODS

### Yeast strains

Yeast strains used in this study are listed in Table S1. Standard culture media were used in this study (Guthrie and Fink, 1991). Conventional gene deletion was performed using a PCR-based method (Longtine *et al*., 1998). Plasmids used for yeast strain construction are listed in Table S2.

### Construction of backbone vectors for genome editing

Oligodeoxyribonucleotides (ODNs) used in this study are listed in Table S3. All ODNs for plasmid construction were purchased from Sigma-Aldrich Japan (Tokyo, Japan) and Eurofins Genomics K.K. (Tokyo, Japan). The backbone vectors for genome editing were constructed using the seamless cloning with HiFi DNA Assembly (E2621, New England Biolabs, Ipswich, MA, USA). Restriction enzymes used for plasmid construction were purchased from New England Biolabs. PCR fragments used for plasmid construction were amplified by Q5 DNA polymerase (M0491, New England Biolabs) or KOD One (KMM-101, TOYOBO, Osaka, Japan) according to the manufacturer’s instructions. *Escherichia coli* competent cells NEB 5-alpha (C2987, New England Biolabs), NEB Stable (C3040, New England Biolabs), or Champion DH5α high (CC5202, SMOBIO Technology, Hsinchu City, Taiwan) were used for transformation to amplify and extract plasmids. Plasmids were extracted by FastGene Plasmid Mini Kit (FG-90502, Nippon Genetics, Tokyo, Japan) or FavorPrep Plasmid Extraction Mini Kit (FAPDE 001-1, FAVORGEN, Ping Tung, Taiwan). Plasmids used in this study are listed in Table S2. The DNA sequence files of the backbone vectors for genome editing are available on our repository at GitHub (https://github.com/poccopen/LOBSTERS).

### Selection of target sequences for genome editing

For designing sgRNAs for SpCas9 and SaCas9, CRISPRdirect (Naito *et al*., 2015) was used to select target sequences. For designing CRISPR RNAs (crRNAs) for enAsCas12a, CRISPOR (Concordet and Haeussler, 2018) was used to select target sequences. The precise positions of target sequences, PAM sites, and cleavage sites for all genome editing experiments are shown in Figure S1.

### Construction of genome editing plasmids

All genome editing plasmids were constructed using the seamless cloning with Golden Gate Assembly using NEB Golden Gate Assembly Kit (BsaI-HF v2) (E1601, New England Biolabs). The ODNs for Golden Gate Assembly were automatically designed with an in-house program.

### Yeast transformation for genome editing

Yeast transformation for the single genome editing (Figure 2) was carried out as described previously (Okada *et al*., 2021). Yeast transformation for the multiple genome editing (Figures 3 and 4) was performed as described below. Yeast cells were cultured overnight in 2 mL of YPAD liquid medium (10 g/L Bacto Yeast Extract, #212750, Thermo Fisher Scientific, Waltham, MA, USA; 20 g/L Bacto Peptone, #211677, Thermo Fisher Scientific; 100 mg/L adenine sulfate, #01990-94, Nacalai tesque, Kyoto, Japan; and 20 g/L glucose, Nacalai tesque) at 25°C with shaking at 250 rpm. The 2-mL culture in the logarithmic growth phase was centrifuged, and the supernatant was removed. The cell pellet was resuspended in 1 mL of 0.1 M lithium acetate solution (#127-01545, FUJIFILM Wako Chemicals, Osaka, Japan). The cell suspension was incubated at 30°C for 45 min. The cell suspension was centrifuged, and the supernatant was removed. The cells were thoroughly mixed with 50 μL of 1 M lithium acetate, 50 μL of 1 M dithiothreitol (#14128-04, Nacalai tesque), 5 μL of Yeastmaker Carrier DNA (10 mg/mL, #630440, Takara Bio, Kusatsu, Japan), 50 μL of DNA solution containing genome editing plasmids (2–6 μg each) and PCR-generated donor fragments for gene fragment insertion (1–10 μg, typically 5 μg each), and 450 μL of polyethylene glycol 4000 (#11574-15, Nacalai tesque). The donor fragments were amplified by Q5 DNA polymerase (New England Biolabs) or KOD One (KMM-101, TOYOBO, Osaka, Japan) according to the manufacturer’s instructions. The samples were incubated at 30°C for 90 min, followed by a 15-min incubation at 42°C. After centrifugation and removal of the supernatant, the cell pellets were resuspended in 50 μL of SC−Ura medium without carbon source (7.4 g/L Yeast nitrogen base without amino acids, #291940, Thermo Fisher Scientific; 855 mg/L CSM−Ura powder, DCS0161, FORMEDIUM, Hunstanton, UK; and 111 mg/L adenine sulfate, Nacalai tesque) and spread on a SCGal−Leu−Ura (Figure 3) or SCGal−His−Leu−Ura (Figure 4) agar plate. The plates were incubated at 30°C for 4–11 days. The colonies were picked and streaked as patches on SCGal−Leu−Ura or SCGal−His−Leu−Ura agar plates and then incubated at 30°C for 1–2 days, followed by colony PCR to check successful genome editing. Target sequences and PAM sites for all experiments are shown in Figure S1. Colony PCR was performed using Q5 DNA polymerase (New England Biolabs) or KOD One (TOYOBO) according to the manufacturer’s instructions. The template DNA solution for colony PCR was prepared by the GC preps method (Blount *et al*., 2016). For gene fragment insertions (Figure 3), successful editing was confirmed by the increased size of PCR products reflecting the insertion of fluorescent protein genes. For nucleotide substitutions (Figure 4), PCR products were purified and subjected to Sanger sequencing to confirm the intended mutations. The positive clones were cultured overnight in 2 mL of YPAD liquid medium. An aliquot (10 μL) of the overnight culture was spotted and streaked on a YPAD agar plate for single colony isolation (30°C for 2 days). Single colonies were picked and streaked on a YPAD agar plate, a SCDex−His agar plate, a SCDex−Leu agar plate, and a SCDex−Ura agar plate to check the loss of the genome-editing plasmids. The Leu^−^ Ura^−^ clones (Figure 3) and the His^−^ Leu^−^ Ura^−^ clones (Figure 4) were re-examined by colony PCR to be successfully genome-edited.

### Yeast colony color assay

The cells after transformation using auxotrophic marker genes (*HIS3, TRP1, LEU2*, and *URA3*) were spread on adenine-limited SCGal agar plates containing 10 mg/L adenine sulfate and lacking the cognate nucleobase or amino acid (uracil, leucine, or histidine, respectively). The cells after transformation using antifungal drug resistance genes (*KanMX, HphMX*, or *NatMX*) were spread on YPD agar plates (10 g/L Bacto Yeast Extract, #212750, Thermo Fisher Scientific, Waltham, MA, USA; 20 g/L Bacto Peptone, #211677, Thermo Fisher Scientific; 20 g/L glucose, Nacalai tesque; and 20 g/L agar, #010-08725, FUJIFILM Wako Chemicals) containing one of the antifungal drugs (200 mg/L G418 disulfate, 300 mg/L hygromycin, or 100 mg/L nourseothricin). The agar plates were incubated at 30°C for 4–5 days, followed by incubation at 4°C for 3–7 days to obtain better coloration of yeast colonies. The images of colonies were captured by a digital camera (TG-6, Olympus, Tokyo, Japan). Editing efficiency was determined by scoring colony color. For each sample, a minimum of 101 colonies and up to 1503 colonies were counted.

### Fluorescence microscopy and image processing

Image acquisition of yeast cells was performed on a microscope (Ti-E, Nikon, Tokyo, Japan) with a 100× objective lens (CFI Apo TIRF 100XC Oil, MRD01991, Nikon), a sCMOS camera (ORCA-Fusion BT, C15440-20UP, Hamamatsu Photonics, Hamamatsu, Japan), and a solid-state illumination light source (SOLA SE II, Lumencor, Beaverton, OR, USA). Image acquisition was controlled by NIS-Elements version 5.3 (Nikon). The binning mode of the camera was set at 2×2 (0.13 μm/pixel). Z-stacks were 13×0.3 μm. For imaging of Cse4-mNeonGreen, a filter set (LED-YFP-A, Semrock, Rochester, NY, USA) was used with excitation light power set at 20% and the exposure time set at 200 msec/frame. For imaging of Cdc3-miRFP682, a filter set (CY5.5-C, Semrock) was used with excitation light power set at 30% and exposure time set at 300 msec/frame. For DIC (differential interference contrast) image acquisition, the exposure time was set at 20 msec/frame. DIC images were captured only at the middle position of the Z-stacks.

Image processing and analysis were performed using Fiji (Schindelin *et al*., 2012). To generate 2-dimensional images of fluorescence channels from the Z-stacks, background subtraction (sliding paraboloid radius set at 10 pixels with disabled smoothing) and maximum projection using 13 Z-slices were performed. Maximum projected fluorescence images and corresponding smoothed DIC images were superimposed. After global adjusting of brightness and contrast and cropping of the images, sequences of representative images were generated.

### Sporulation assay

Yeast cells were cultured in 2 mL of YPAc liquid medium (10 g/L Bacto Yeast Extract, Thermo Fisher Scientific; 20 g/L Bacto Peptone, Thermo Fisher Scientific;100 mg/L adenine sulfate, Nacalai tesque; and 20 g/L potassium acetate, #28405-05, Nacalai tesque) for 2 days at 30°C with shaking at 250 rpm. Then the cells were washed with the sporulation medium (20 g/L potassium acetate, Nacalai tesque) twice, and the cells were resuspended in 2 mL of the sporulation medium, followed by shaking at 250 rpm at 30°C for 2 days. The cells were centrifuged, and the supernatant was removed. The cells were fixed with 70% ethanol for 45 min at room temperature. The fixed cells were washed with PBS twice and resuspended in PBS. DIC images of the cells were acquired on a microscope (Ti-E, Nikon) with a 100× objective lens (CFI Apo TIRF 100XC Oil, Nikon), a sCMOS camera (ORCA-Fusion BT, Hamamatsu Photonics), and a solid-state illumination light source (SOLA SE II, Lumencor). Image acquisition was controlled by NIS-Elements version 5.3 (Nikon). For each strain, sporulation efficiency was determined by counting 61–185 cells per sample.

## Abbreviations

Cas: CRISPR-associated protein;
CRISPR: clustered regularly interspaced short palindromic repeats;
crRNA: CRISPR RNA;
DIC: differential interference contrast;
LOBSTERS: League Of Backbone plasmid vector Series To Expand the Range of Selection markers;
ORF: open reading frame;
PAM: protospacer adjacent motif;
PCR: polymerase chain reaction;
QTV: quantitative trait variant;
sgRNA: single guide RNA.

## DATA AVAILABILITY

All 21 LOBSTERS backbone plasmids will be available from NBRP Yeast Resource Center (https://yeast.nig.ac.jp/yeast/). Plasmid sequence files and the source codes of programs for ODN design are available from our repository at GitHub (https://github.com/poccopen/LOBSTERS). Other strains and plasmids are available upon request. The authors state that all data necessary for confirming the conclusions presented here are represented fully within the article.

## ACKNOWLEDGMENTS

Yeast strains (BY5445, BY5446, BY19600, and W303-1A) were provided by the National Bio-Resource Project (NBRP), Japan. We thank Hiroaki Takesue, Suchin Towa, and Yuki Sugiyama for their pilot use of the genome editing systems during the initial phase of the project. This work was supported by JST CREST Grant Number JPMJCR19S1 and JSPS KAKENHI Grant Numbers 24H00505 and 24K02015.

