## Supplemental Figure S1 for "LOBSTERS: a modular vector series with a diverse set of Cas nucleases and plasmid selection markers for multiplex genome editing in *Saccharomyces cerevisiae*"

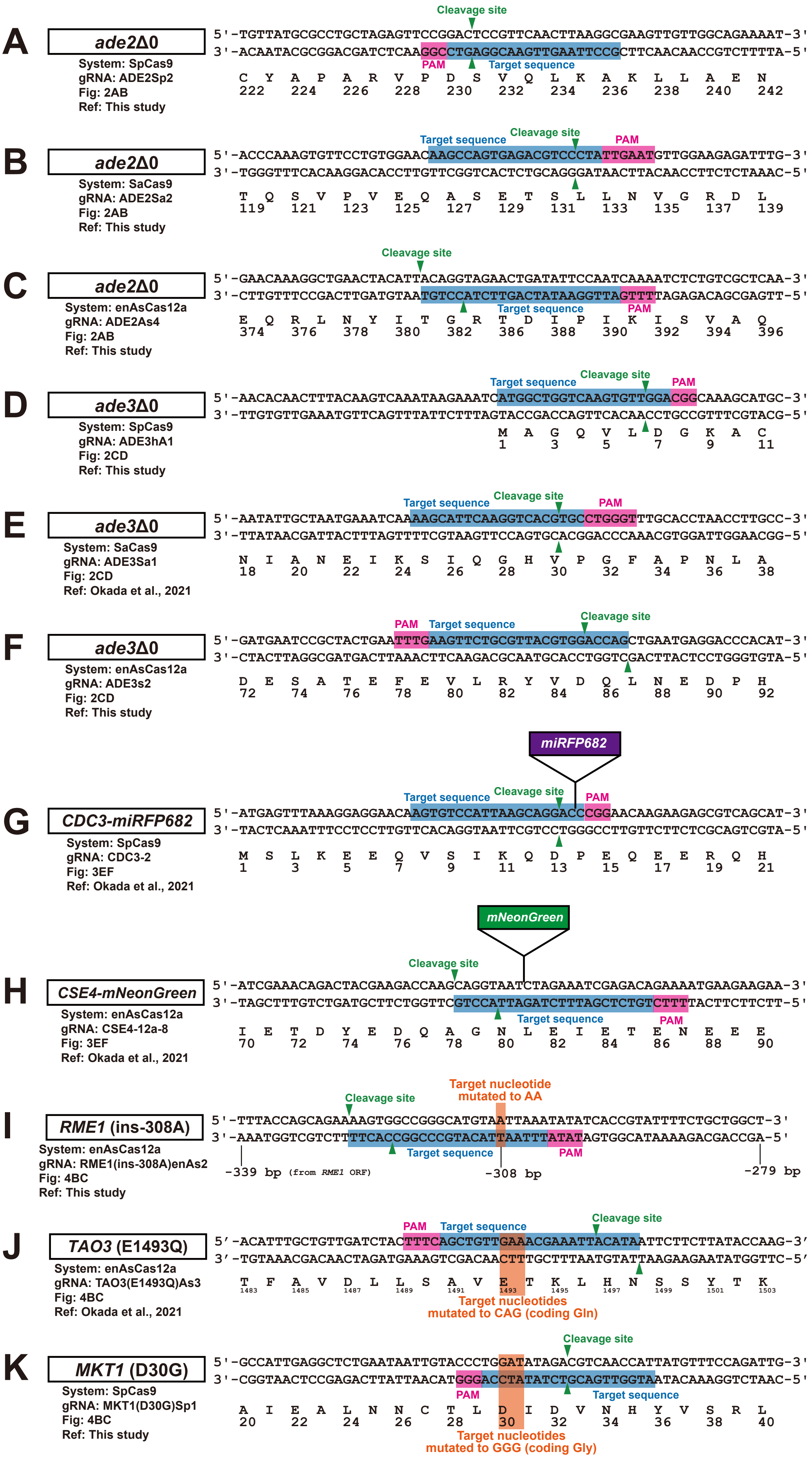


*(legend on next page)*

(legend on next page)

**Figure S1.** Target sequences, PAM sites, and cleavage positions for genome editing experiments. (A-C) Target sequences for complete deletion of the *ADE2* ORF using SpCas9, SaCas9, and enAsCas12a systems, respectively. (D-F) Target sequences for complete deletion of the *ADE3* ORF using SpCas9, SaCas9, and enAsCas12a systems, respectively. (G) Target sequence for *CDC3-miRFP682* gene fragment insertion using SpCas9. (H) Target sequence for *CSE4-mNeonGreen* gene fragment insertion using enAsCas12a. (I-K) Target sequences for introducing quantitative trait nucleotides: *RME1*(ins-308A) using enAsCas12a, *TAO3*(E1493Q) using enAsCas12a, and *MKT1* (D30G) using SpCas9, respectively. For each target, the top strand (5' to 3') and bottom strand (3' to 5') of genomic DNA sequences are shown with the corresponding amino acid sequence below (single-letter code). Numbers below amino acids indicate the position from the translation start site or relevant genomic landmark. PAM sequences are highlighted in pink boxes. Target sequences recognized by guide RNAs are highlighted in blue boxes (for SpCas9 and SaCas9) or green boxes (for enAsCas12a). Cleavage sites are indicated by arrows. Nucleotides to be mutated or inserted are shown in red text. For panel I (*RME1*), the position relative to the *RME1* ORF start site is indicated.
