## Supplemental Tables S1-S3 for "LOBSTERS: a modular vector series with a diverse set of Cas nucleases and plasmid selection markers for multiplex genome editing in *Saccharomyces cerevisiae*"

**Table S1. Yeast strains used in this study.**

| Strain name | Strain number | MAT | Relevant genotype | Strain background | Relevant figures | sgRNAs/crRNAs | Reference | Source |
|---|---|---|---|---|---|---|---|---|
| BY4741 | YIT5001 | a | *his3Δ1 leu2Δ0 met15Δ0 ura3Δ0* | BY4741 | 2B, 4CDE | - | (Brachmann et al., 1998) | |
| BY4742 | YIT5002 | alpha | *his3Δ1 leu2Δ0 lys2Δ0 ura3Δ0* | BY4742 | 3EGF, 4CDE | - | (Brachmann et al., 1998) | |
| YIT6460 | YIT6460 | a | *ade2Δ::HIS3MX6* | BY4741 | 2D | - | (Okada et al., 2021) | |
| W303-1A | YIT5296 | a | *ade2-1 can1-100 his3-11,15 leu2-3,112 trp1-1 ura3-1 rad5-535* | W303 | 2D | - | (Thomas & Rothstein, 1989) | http://yeast.lab.nig.ac.jp/nig/ |
| YIT11433 | YIT11433 | alpha | *CDC3-ymiRFP682(int-aa13)* | BY4742 | 3E | CDC3-2 | This study | |
| YIT11435 | YIT11435 | alpha | *CDC3-ymiRFP682(int-aa13)* | BY4742 | 3E | CDC3-2 | This study | |
| YIT11436 | YIT11436 | alpha | *CDC3-ymiRFP682(int-aa13)* | BY4742 | 3E | CDC3-2 | This study | |
| YIT11434 | YIT11434 | alpha | *CSE4-ymNeonGreen(int-aa80)* | BY4742 | 3E | CSE4-12a-8 | This study | |
| YIT11534 | YIT11534 | alpha | *CDC3-ymiRFP682(int-aa13) CSE4-ymNeonGreen(int-aa80)* | BY4742 | 3FG | CDC3-2, CSE4-12a-8 | This study | |
| BY5445 | YIT5164 | a | *ura3 leu2::hisG trp1::hisG lys2 ho::LYS2* | SK1 | 4DE | - | (Alani et al., 1987) | http://yeast.lab.nig.ac.jp/nig/ |
| BY5446 | YIT5165 | alpha | *ura3 leu2::hisG trp1::hisG lys2 ho::LYS2* | SK1 | 4DE | - | (Alani et al., 1987) | http://yeast.lab.nig.ac.jp/nig/ |
| BY19600 | YIT5166 | a/alpha | *ura3/ura3 leu2::hisG/leu2::hisG trp1::hisG/trp1::hisG lys2/lys2 ho::LYS2/ho::LYS2* | SK1 | 4CDE | - | (Alani et al., 1987) | http://yeast.lab.nig.ac.jp/nig/ |
| YIT11649 | YIT11649 | a/alpha | *his3Δ1*/his3Δ1 *leu2Δ0*/leu2Δ0 *met15Δ0*/MET15 *LYS2/lys2Δ0 ura3Δ0*/ura3Δ0 | BY4741 x BY4742 | 4DE | - | This study | |
| YIT12052 | YIT12052 | a | *RME1(ins-308A) TAO3(E1493Q) MKT1(D30G)* | BY4741 | 4CDE | RME1(ins-308A)enAs2, TAO3(E1493Q)As3, MKT1(D30G)Sp1 | This study | |
| YIT12053 | YIT12053 | alpha | *RME1(ins-308A) TAO3(E1493Q) MKT1(D30G)* | BY4742 | 4CDE | RME1(ins-308A)enAs2, TAO3(E1493Q)As3, MKT1(D30G)Sp1 | This study | |
| YIT12075 | YIT12075 | a/alpha | *RME1(ins-308A)/RME1(ins-308A) TAO3(E1493Q)/TAO3(E1493Q) MKT1(D30G)/MKT1(D30G)* | BY4741 x BY4742 | 4CDE | RME1(ins-308A)enAs2, TAO3(E1493Q)As3, MKT1(D30G)Sp1 | This study | |
| YIT12076 | YIT12076 | a/alpha | *RME1(ins-308A)/RME1(ins-308A) TAO3(E1493Q)/TAO3(E1493Q) MKT1(D30G)/MKT1(D30G)* | BY4741 x BY4742 | 4CDE | RME1(ins-308A)enAs2, TAO3(E1493Q)As3, MKT1(D30G)Sp1 | This study | |
| YIT12077 | YIT12077 | a/alpha | *RME1(ins-308A)/RME1(ins-308A) TAO3(E1493Q)/TAO3(E1493Q) MKT1(D30G)/MKT1(D30G)* | BY4741 x BY4742 | 4CDE | RME1(ins-308A)enAs2, TAO3(E1493Q)As3, MKT1(D30G)Sp1 | This study | |
| YIT12078 | YIT12078 | a/alpha | *RME1(ins-308A)/RME1(ins-308A) TAO3(E1493Q)/TAO3(E1493Q) MKT1(D30G)/MKT1(D30G)* | BY4741 x BY4742 | 4CDE | RME1(ins-308A)enAs2, TAO3(E1493Q)As3, MKT1(D30G)Sp1 | This study | |

**Table S2. Plasmids used in this study.**

| Plasmid number | Plasmid name | Plasmid name (alias) | Relevant figures | Reference |
|---|---|---|---|---|
| 16-15 | pGAL1-Cas9-tADH1-pGAL1-2BsaI-sgRNAFE(empty)-HDV-tCYC1-CU | pLOBSTER-SpCas9-Ura | 1A | (Okada et al., 2021) |
| 25-31 | pGAL1-Cas9-tADH1-pGAL1-2Bsa-sgRNAFE(empty)-HDV-tCYC1-CLeu | pLOBSTER-SpCas9-Leu | 1A | This study |
| 25-30 | pGAL1-Cas9-tADH1-pGAL1-2Bsa-sgRNAFE(empty)-HDV-tCYC1-CHis | pLOBSTER-SpCas9-His | 1A | This study |
| 25-27 | pGAL1-Cas9-tADH1-pGAL1-2Bsa-sgRNAFE(empty)-HDV-tCYC1-CKan | pLOBSTER-SpCas9-Kan | 1A | This study |
| 25-28 | pGAL1-Cas9-tADH1-pGAL1-2Bsa-sgRNAFE(empty)-HDV-tCYC1-CHyg | pLOBSTER-SpCas9-Hyg | 1A | This study |
| 25-29 | pGAL1-Cas9-tADH1-pGAL1-2Bsa-sgRNAFE(empty)-HDV-tCYC1-CNat | pLOBSTER-SpCas9-Nat | 1A | This study |
| 39-46 | pGAL1-Cas9-tADH1-pGAL1-2Bsa-sgRNAFE(empty)-HDV-tCYC1-CTrp | pLOBSTER-SpCas9-Trp | 1A | This study |
| 17-31 | pGAL1-ySaCas9-tADH1-pGAL1-2BsaI-SagRNA(empty)-HDV-tCYC1-CU | pLOBSTER-SaCas9-Ura | 1B | (Okada et al., 2021) |
| 25-36 | pGAL1-ySaCas9-tADH1-pGAL1-2Bsa-SagRNA(empty)-HDV-tCYC1-CLeu | pLOBSTER-SaCas9-Leu | 1B | This study |
| 25-35 | pGAL1-ySaCas9-tADH1-pGAL1-2Bsa-SagRNA(empty)-HDV-tCYC1-CHis | pLOBSTER-SaCas9-His | 1B | This study |
| 25-32 | pGAL1-ySaCas9-tADH1-pGAL1-2Bsa-SagRNA(empty)-HDV-tCYC1-CKan | pLOBSTER-SaCas9-Kan | 1B | This study |
| 25-33 | pGAL1-ySaCas9-tADH1-pGAL1-2Bsa-SagRNA(empty)-HDV-tCYC1-CHyg | pLOBSTER-SaCas9-Hyg | 1B | This study |
| 25-34 | pGAL1-ySaCas9-tADH1-pGAL1-2Bsa-SagRNA(empty)-HDV-tCYC1-CNat | pLOBSTER-SaCas9-Nat | 1B | This study |
| 39-48 | pGAL1-ySaCas9-tADH1-pGAL1-2Bsa-SagRNA(empty)-HDV-tCYC1-CTrp | pLOBSTER-SaCas9-Trp | 1B | This study |
| 16-16 | pGAL1-yenAsCas12a-tADH1-pGAL1-tRNA(Gly)-HindIII-AsCpf1gRNA(empty)-2BsaI-U4AU4-HDV-tCYC1-CU | pLOBSTER-enAsCas12a-Ura | 1C | (Okada et al., 2021) |
| 25-41 | pGAL1-yenAsCas12a-tADH1-pGAL1-tRNA(Gly)-AscrRNA(empty)-2Bsa-U4AU4-HDV-tCYC1-CLeu | pLOBSTER-enAsCas12a-Leu | 1C | This study |
| 25-40 | pGAL1-yenAsCas12a-tADH1-pGAL1-tRNA(Gly)-AscrRNA(empty)-2Bsa-U4AU4-HDV-tCYC1-CHis | pLOBSTER-enAsCas12a-His | 1C | This study |
| 25-37 | pGAL1-yenAsCas12a-tADH1-pGAL1-tRNA(Gly)-AscrRNA(empty)-2Bsa-U4AU4-HDV-tCYC1-CKan | pLOBSTER-enAsCas12a-Kan | 1C | This study |
| 25-38 | pGAL1-yenAsCas12a-tADH1-pGAL1-tRNA(Gly)-AscrRNA(empty)-2Bsa-U4AU4-HDV-tCYC1-CHyg | pLOBSTER-enAsCas12a-Hyg | 1C | This study |
| 25-39 | pGAL1-yenAsCas12a-tADH1-pGAL1-tRNA(Gly)-AscrRNA(empty)-2Bsa-U4AU4-HDV-tCYC1-CNat | pLOBSTER-enAsCas12a-Nat | 1C | This study |
| 39-47 | pGAL1-yenAsCas12a-tADH1-pGAL1-tRNA(Gly)-AscrRNA(empty)-2Bsa-U4AU4-HDV-tCYC1-CTrp | pLOBSTER-enAsCas12a-Trp | 1C | This study |
| 1-50 | YCplac33 | YCpURA3 | 2B | (Gietz and Sugino, 1988) |
| 1-51 | YCplac111 | YCpLEU2 | 2B | (Gietz and Sugino, 1988) |
| 32-26 | YCpHIS3 | - | 2B | This study |
| 33-49 | pGAL1-Cas9-tADH1-pGAL1-HH-sgRNAFE(ADE2Sp2)-HDV-tCYC-CUra | pLOBSTER-SpCas9-Ura(ADE2Sp2) | 2B | This study |
| 32-79 | pGAL1-Cas9-tADH1-pGAL1-HH-sgRNAFE(ADE2Sp2)-HDV-tCYC-CLeu | pLOBSTER-SpCas9-Leu(ADE2Sp2) | 2B | This study |
| 32-67 | pGAL1-Cas9-tADH1-pGAL1-HH-sgRNAFE(ADE2Sp2)-HDV-tCYC-CHis | pLOBSTER-SpCas9-His(ADE2Sp2) | 2B | This study |
| 33-51 | pGAL1-ySaCas9-tADH1-pGAL1-HH-SagRNA(ADE2Sa2)-HDV-tCYC-CUra | pLOBSTER-SaCas9-Ura(ADE2Sa2) | 2B | This study |
| 33-2 | pGAL1-ySaCas9-tADH1-pGAL1-HH-SagRNA(ADE2Sa2)-HDV-tCYC-CLeu | pLOBSTER-SaCas9-Leu(ADE2Sa2) | 2B | This study |
| 32-71 | pGAL1-ySaCas9-tADH1-pGAL1-HH-SagRNA(ADE2Sa2)-HDV-tCYC-CHis | pLOBSTER-SaCas9-His(ADE2Sa2) | 2B | This study |
| 33-53 | pGAL1-yenAsCas12a-tADH1-pGAL1-tRNA(Gly)-AscrRNA(ADE2As4)-U4AU4-HDV-tCYC1-CUra | pLOBSTER-enAsCas12a-Ura(ADE2As4) | 2B | This study |
| 33-8 | pGAL1-yenAsCas12a-tADH1-pGAL1-tRNA(Gly)-AscrRNA(ADE2As4)-U4AU4-HDV-tCYC1-CLeu | pLOBSTER-enAsCas12a-Leu(ADE2As4) | 2B | This study |
| 32-77 | pGAL1-yenAsCas12a-tADH1-pGAL1-tRNA(Gly)-AscrRNA(ADE2As4)-U4AU4-HDV-tCYC1-CHis | pLOBSTER-enAsCas12a-His(ADE2As4) | 2B | This study |
| 23-59 | YCp-KanMX | - | 2D | This study |
| 23-60 | YCp-HphMX | - | 2D | This study |
| 23-61 | YCp-NatMX | - | 2D | This study |
| 1-47 | YCplac22 | YCpTRP1 | 2D | (Gietz and Sugino, 1988) |
| 28-8 | pGAL1-Cas9-tADH1-pGAL1-HH-sgRNAFE(ADE3hA1)-HDV-tCYC1-CKan | pLOBSTER-SpCas9-Kan(ADE3hA1) | 2D | This study |
| 28-9 | pGAL1-Cas9-tADH1-pGAL1-HH-sgRNAFE(ADE3hA1)-HDV-tCYC1-CHyg | pLOBSTER-SpCas9-Hyg(ADE3hA1) | 2D | This study |
| 28-10 | pGAL1-Cas9-tADH1-pGAL1-HH-sgRNAFE(ADE3hA1)-HDV-tCYC1-CNat | pLOBSTER-SpCas9-Nat(ADE3hA1) | 2D | This study |
| 39-66 | pGAL1-Cas9-tADH1-pGAL1-HH-sgRNAFE(ADE3hA1)-HDV-tCYC1-CTrp | pLOBSTER-SpCas9-Trp(ADE3hA1) | 2D | This study |
| 28-3 | pGAL1-ySaCas9-tADH1-pGAL1-HH-SagRNA(ADE3Sa1)-HDV-tCYC1-CKan | pLOBSTER-SaCas9-Kan(ADE3Sa1) | 2D | This study |
| 28-4 | pGAL1-ySaCas9-tADH1-pGAL1-HH-SagRNA(ADE3Sa1)-HDV-tCYC1-CHyg | pLOBSTER-SaCas9-Hyg(ADE3Sa1) | 2D | This study |
| 28-5 | pGAL1-ySaCas9-tADH1-pGAL1-HH-SagRNA(ADE3Sa1)-HDV-tCYC1-CNat | pLOBSTER-SaCas9-Nat(ADE3Sa1) | 2D | This study |
| 39-68 | pGAL1-ySaCas9-tADH1-pGAL1-HH-SagRNA(ADE3Sa1)-HDV-tCYC1-CTrp | pLOBSTER-SaCas9-Trp(ADE3Sa1) | 2D | This study |
| 28-26 | pGAL1-yenAsCas12a-tADH1-pGAL1-tRNA(Gly)-AscrRNA(ADE3s2)-U4AU4-HDV-tCYC1-CKan | pLOBSTER-enAsCas12a-Kan(ADE3s2) | 2D | This study |
| 28-27 | pGAL1-yenAsCas12a-tADH1-pGAL1-tRNA(Gly)-AscrRNA(ADE3s2)-U4AU4-HDV-tCYC1-CHyg | pLOBSTER-enAsCas12a-Hyg(ADE3s2) | 2D | This study |
| 28-28 | pGAL1-yenAsCas12a-tADH1-pGAL1-tRNA(Gly)-AscrRNA(ADE3s2)-U4AU4-HDV-tCYC1-CNat | pLOBSTER-enAsCas12a-Nat(ADE3s2) | 2D | This study |
| 39-67 | pGAL1-yenAsCas12a-tADH1-pGAL1-tRNA(Gly)-AscrRNA(ADE3s2)-U4AU4-HDV-tCYC1-CTrp | pLOBSTER-enAsCas12a-Trp(ADE3s2) | 2D | This study |
| 27-77 | pGAL1-Cas9-tADH1-pGAL1-HH-sgRNAFE(CDC3-2)-HDV-tCYC1-CU | pLOBSTER-SpCas9-Ura(CDC3-2) | 3EF | This study |
| 28-49 | pGAL1-yenAsCas12a-tADH1-pGAL1-tRNA(Gly)-AscrRNA(CSE4-12a-8)-U4AU4-HDV-tCYC1-CLeu | pLOBSTER-enAsCas12a-Leu(CSE4-12a-8) | 3EF | This study |
| 29-43 | pGAL1-yenAsCas12a-tADH1-pGAL1-tRNA(Gly)-AscrRNA(RME1(ins-308A)enAs2)-U4AU4-HDV-tCYC1-CLeu | pLOBSTER-enAsCas12a-Leu(RME1(ins-308A)enAs2) | 4BC | This study |
| 24-54 | pGAL1-yenAsCas12a-tADH1-pGAL1-tRNA(Gly)-AscrRNA(TAO3(E1493Q)As3)-U4AU4-HDV-tCYC1-CU | pLOBSTER-enAsCas12a-Ura(TAO3(E1493Q)As3) | 4BC | (Okada et al., 2021) |
| 29-44 | pGAL1-Cas9-tADH1-pGAL1-HH-sgRNAFE(MKT1(D30G)Sp1)-HDV-tCYC1-CHis | pLOBSTER-SpCas9-His(MKT1(D30G)) | 4BC | This study |

**Table S3. Oligo DNA sequences used in this study.**

| Oligo DNA number | Oligo DNA name | Oligo DNA sequence | Cas | Description | Relevant figures |
|---|---|---|---|---|---|
| 87-25 | ADE2Sp2F | GGAGTAAGGCCTGATGAGTCCGTGAGGACGAAACGAGTAAGCTCGTCGCCTTAAGTTGAACGGAGTC | SpCas9 | Genome-editing plasmid construction | 2B |
| 87-37 | ADE2Sp2R | AAACGACTCCGTTCAACTTAAGGCGACGAGCTTACTCGTTTCGTCCTCACGGACTCATCAGGCCTTA | SpCas9 | Genome-editing plasmid construction | 2B |
| 87-29 | ADE2Sa2F | GGAGTGGCTTCTGATGAGTCCGTGAGGACGAAACGAGTAAGCTCGTCAAGCCAGTGAGACGTCCCTA | SaCas9 | Genome-editing plasmid construction | 2B |
| 87-41 | ADE2Sa2R | TAACTAGGGACGTCTCACTGGCTTGACGAGCTTACTCGTTTCGTCCTCACGGACTCATCAGAAGCCA | SaCas9 | Genome-editing plasmid construction | 2B |
| 87-35 | ADE2As4F | AGATATTGGAATATCAGTTCTACCTGT | enAsCas12a | Genome-editing plasmid construction | 2B |
| 87-47 | ADE2As4R | AAAAACAGGTAGAACTGATATTCCAAT | enAsCas12a | Genome-editing plasmid construction | 2B |
| 88-26 | ADE2up50down50-F | CCTACTATAACAATCAAGAAAAACAAGAAAATCGGACAAAACAATCAAGTTATATAAGTTTATTG | - | Preparation of donor fragment for ADE2 deletion | 2B |
| 88-27 | ADE2up50down50-R | ATATCATTTTATAATTATTTGCTGTACAAGTATATCAATAAACTTATATAACTTGATTGTTTTGT | - | Preparation of donor fragment for ADE2 deletion | 2B |
| 84-39 | ADE2-87-5 | GGTGCGTAAAATCGTTGGAT | - | ADE2 deletion PCR check | 2B |
| 84-40 | ADE2+100-3 | TGTATGAAGTCCACATTTGATGTAA | - | ADE2 deletion PCR check | 2B |
| 40-73 | ADE3hA1-F | GGAGAGCCATCTGATGAGTCCGTGAGGACGAAACGAGTAAGCTCGTCATGGCTGGTCAAGTGTTGGA | SpCas9 | Genome-editing plasmid construction | 2D |
| 40-81 | ADE3hA1-R | AAACTCCAACACTTGACCAGCCATGACGAGCTTACTCGTTTCGTCCTCACGGACTCATCAGATGGCT | SpCas9 | Genome-editing plasmid construction | 2D |
| 45-33 | ADE3-Sa1-F | GGAGATGCTTCTGATGAGTCCGTGAGGACGAAACGAGTAAGCTCGTCAAGCATTCAAGGTCACGTGC | SaCas9 | Genome-editing plasmid construction | 2D |
| 45-37 | ADE3-Sa1-R | TAACGCACGTGACCTTGAATGCTTGACGAGCTTACTCGTTTCGTCCTCACGGACTCATCAGAAGCAT | SaCas9 | Genome-editing plasmid construction | 2D |
| 71-25 | ADE3s2F | agatAAGTTCTGCGTTACGTGGACCAG | enAsCas12a | Genome-editing plasmid construction | 2D |
| 71-26 | ADE3s2R | aaaaCTGGTCCACGTAACGCAGAACTT | enAsCas12a | Genome-editing plasmid construction | 2D |
| 76-12 | ADE3updown50-F | TGAGACCAGGTAACGAGACGAACACAACTTTACAAGTCAAATAAGAAATCGACAACTTAATTATA | - | Preparation of donor fragment for ADE3 deletion | 2D |
| 76-13 | ADE3updown50-R | AAAAAAACTTTTGCATTTGTCTTTATTAAATTCTATATAATTAAGTTGTCGATTTCTTATTTGAC | - | Preparation of donor fragment for ADE3 deletion | 2D |
| 46-60 | ADE3-221-5 | ATTTGCCATGACTCCTCCCA | - | ADE3 deletion PCR check | 2D |
| 46-61 | ADE3+257-3 | CTCATTAATGCGTCTCCCCG | - | ADE3 deletion PCR check | 2D |
| 70-22 | CDC3-2Fv2 | GGAGGACACTCTGATGAGTCCGTGAGGACGAAACGAGTAAGCTCGTCAGTGTCCATTAAGCAGGACC | SpCas9 | Genome-editing plasmid construction | 3FG |
| 70-26 | CDC3-2Rv2 | AAACGGTCCTGCTTAATGGACACTGACGAGCTTACTCGTTTCGTCCTCACGGACTCATCAGAGTGTC | SpCas9 | Genome-editing plasmid construction | 3FG |
| 73-37 | CDC3-ymiRFP682-13F | GTTGCCATGAGTTTAAAGGAGGAACAAGTGTCCATTAAGCAGGACATGGCTGAAGGCAGCGTCGC | - | Preparation of donor fragment for genome-editing | 3FG |
| 73-38 | CDC3-ymiRFP682-13R | CTGAACATCATTGAATTGATCATGCTGACGCTCTTCTTGTTCCGGCGCGCCCTCTTCCATCACTC | - | Preparation of donor fragment for genome-editing | 3FG |
| 38-81 | CDC3-51-5 | TCCAGGATCACACGACAACT | - | CDC3-mScarlet-I PCR check | 3FG |
| 39-1 | CDC3aa38-3 | TCCATCATGATCCTGCGACT | - | CDC3-mScarlet-I PCR check | 3FG |
| 37-67 | CSE4-12a-8F | AGATTGTCTCGATTTCTAGATTACCTG | enAsCas12a | Genome-editing plasmid construction | 3FG |
| 37-75 | CSE4-12a-8R | AAAACAGGTAATCTAGAAATCGAGACA | enAsCas12a | Genome-editing plasmid construction | 3FG |
| 56-34 | CSE4-12a-8Fv2-mNG | AGTGACCTAGATATCGAAACAGACTACGAAGACCAAGCAGGTAATATGGTCTCAAAAGGCGAAGA | - | Preparation of donor fragment for genome-editing | 3FG |
| 56-35 | CSE4-12a-8Rv2-mNG | AGTTTCCATTTCAGCTTCTTCTTCATTTTCTGTCTCGATTTCTAGCTTGTAAAGTTCGTCCATGC | - | Preparation of donor fragment for genome-editing | 3FG |
| 38-56 | CSE4-175-5 | ATCCCACTGTGTCGCGAAAT | - | CSE4-mNeonGreen PCR check | 3FG |
| 38-57 | CSE4aa186-3 | ACGCCATAATCGCCATTGAC | - | CSE4-mNeonGreen PCR check | 3FG |
| 62-48 | RME1(ins-308A)enAs2-F | AGATTTTAATACATGCCCGGCCACTTT | enAsCas12a | Genome-editing plasmid construction | 4BC |
| 62-52 | RME1(ins-308A)enAs2-R | AAAAAAAGTGGCCGGGCATGTATTAAA | enAsCas12a | Genome-editing plasmid construction | 4BC |
| 62-74 | RME1(ins-308A)-Donor-F | ATAAGAAAATCCGAACTTTTCCGGTGGGCTTTACCAGCAGAAAAGTGGCCGGGCATGTAaTTAAATATAT | - | Preparation of donor fragment for genome-editing | 4BC |
| 62-75 | RME1(ins-308A)-Donor-R | TCTGGCCTTTGTTCTCTGCATAAGAAAAAGCCAGCAGAAAATACGGTGATATATTTAAtTACATGCCCGG | - | Preparation of donor fragment for genome-editing | 4BC |
| 63-57 | RME1-493-5 | TGCCAGAAAGAAACCGCAAA | - | RME1 Sanger sequencing | 4BC |
| 63-58 | RME1-67-3 | ACAATGTCAGTTCCAATGCGT | - | RME1 Sanger sequencing | 4BC |
| 62-29 | TAO3(E1493Q)As3-F | AGATAGCTGTTGAAACGAAATTACATA | enAsCas12a | Genome-editing plasmid construction | 4BC |
| 62-32 | TAO3(E1493Q)As3-R | AAAATATGTAATTTCGTTTCAACAGCT | enAsCas12a | Genome-editing plasmid construction | 4BC |
| 62-33 | TAO3(E1493Q)-Donor-F | TTTAGTGGCAGATAAGGAAGATACAAGGACATTTGCTGTTGATCTACTTTCAGCTGTTcagACGAAATTA | - | Preparation of donor fragment for genome-editing | 4BC |
| 62-34 | TAO3(E1493Q)-Donor-R | AAGAATTTGCAAGTCTCTCCTTAAATACCTTGGTATAAGAAGAATTATGTAATTTCGTctgAACAGCTGA | - | Preparation of donor fragment for genome-editing | 4BC |
| 63-48 | TAO3aa1371-5 | TGGACGATGATTTAGAGTTGCC | - | TAO3 Sanger sequencing | 4BC |
| 63-50 | TAO3aa1567-3 | CCCAAGGTACCATAAGCACC | - | TAO3 Sanger sequencing | 4BC |
| 62-36 | MKT1(D30G)Sp1-F | GGAGAACCATCTGATGAGTCCGTGAGGACGAAACGAGTAAGCTCGTCATGGTTGACGTCTATATCCA | SpCas9 | Genome-editing plasmid construction | 4BC |
| 62-39 | MKT1(D30G)Sp1-R | AAACTGGATATAGACGTCAACCATGACGAGCTTACTCGTTTCGTCCTCACGGACTCATCAGATGGTT | SpCas9 | Genome-editing plasmid construction | 4BC |
| 62-78 | MKT1(D30G)-Donor-F | TTTCGAAAGAGGTCTAGTAGGATCCTATGCCATTGAGGCTCTGAATAATTGTACCCTGGggATAGACGTC | - | Preparation of donor fragment for genome-editing | 4BC |
| 62-79 | MKT1(D30G)-Donor-R | CCAAATACTGTTCTCTTTTATTGGTCAACAATCTGGAAACATAATGGTTGACGTCTATccCCAGGGTACA | - | Preparation of donor fragment for genome-editing | 4BC |
| 81-36 | MKT1-212-F | TTTTTGCCTTTCCTTTTAAATTC | - | MKT1 Sanger sequencing | 4BC |
| 81-37 | MKT1aa123-R | CGTGTTGGATCTTGTTGTGG | - | MKT1 Sanger sequencing | 4BC |
